# PyGenePlexus: A Python package for gene discovery using network-based machine learning

**DOI:** 10.1101/2022.07.02.498552

**Authors:** Christopher A Mancuso, Renming Liu, Arjun Krishnan

## Abstract

*PyGenePlexus* is a Python package that enables a user to gain insight into any gene set of interest based on a molecular interaction network using supervised machine learning. *PyGenePlexus* provides predictions of how associated every gene in the network is to the input gene set, offers interpretability by comparing the model trained on the input gene set to models trained on thousands of known gene sets, and returns the network connectivity of the top predicted genes.

**Availability and Implementation:** https://pypi.org/project/geneplexus/ and https://github.com/krishnanlab/PyGenePlexus

## Introduction

Most functions, phenotypes, and diseases are orchestrated by the complex interactions of many genes. To probe these biological contexts, researchers routinely generate sets of genes specific to those contexts using high-throughput, high-coverage technologies (Heller, 2002; Wang *et al*., 2009). Numerous databases contain curated gene sets pertaining to various processes (Ashburner *et al*., 2000; The Gene Ontology Consortium, 2019), diseases (Piñero *et al*., 2015, 2017; Schriml *et al*., 2019) and traits (Choobdar *et al*., 2019) via publicly available databases. However, these gene sets are often incomplete, noisy, and provide no information on how the genes in the set interact with each other, making it hard to fully understand the underlying biology that connects the genes. Hence, developing computational approaches that can provide insights into gene sets is a grand challenge in biomedical research (Yang *et al*., 2011; Piro and Cunto, 2012; Jiang *et al*., 2016).

Computational methods that incorporate information from genome-wide, context-specific molecular-networks have recently shown state-of-the-art results in the task of prioritizing genes of interest and predicting other novel genes that may be highly related to the original gene set (Warde-Farley *et al*., 2010; Köhler *et al*., 2008; Greene *et al*., 2015; Krishnan *et al*., 2016). In a previous work, we have shown that using a supervised machine learning (ML) model that uses the connections from a genome-wide molecular network as the features in a supervised ML model (referred to as *GenePlexus*) is a robust, data-driven way to computationally predict how associated a new gene is to a given input gene set (Liu *et al*., 2020). *GenePlexus* produces more accurate gene classification performance compared to traditional label propagation based methods on diverse sets of tasks including functional, disease, and trait predictions. In this work, we present *PyGenePlexus*, a python package that enables users to easily run the *GenePlexus* method on their input gene sets of choice on the command line [Fig. 1A].

**Figure 1.**
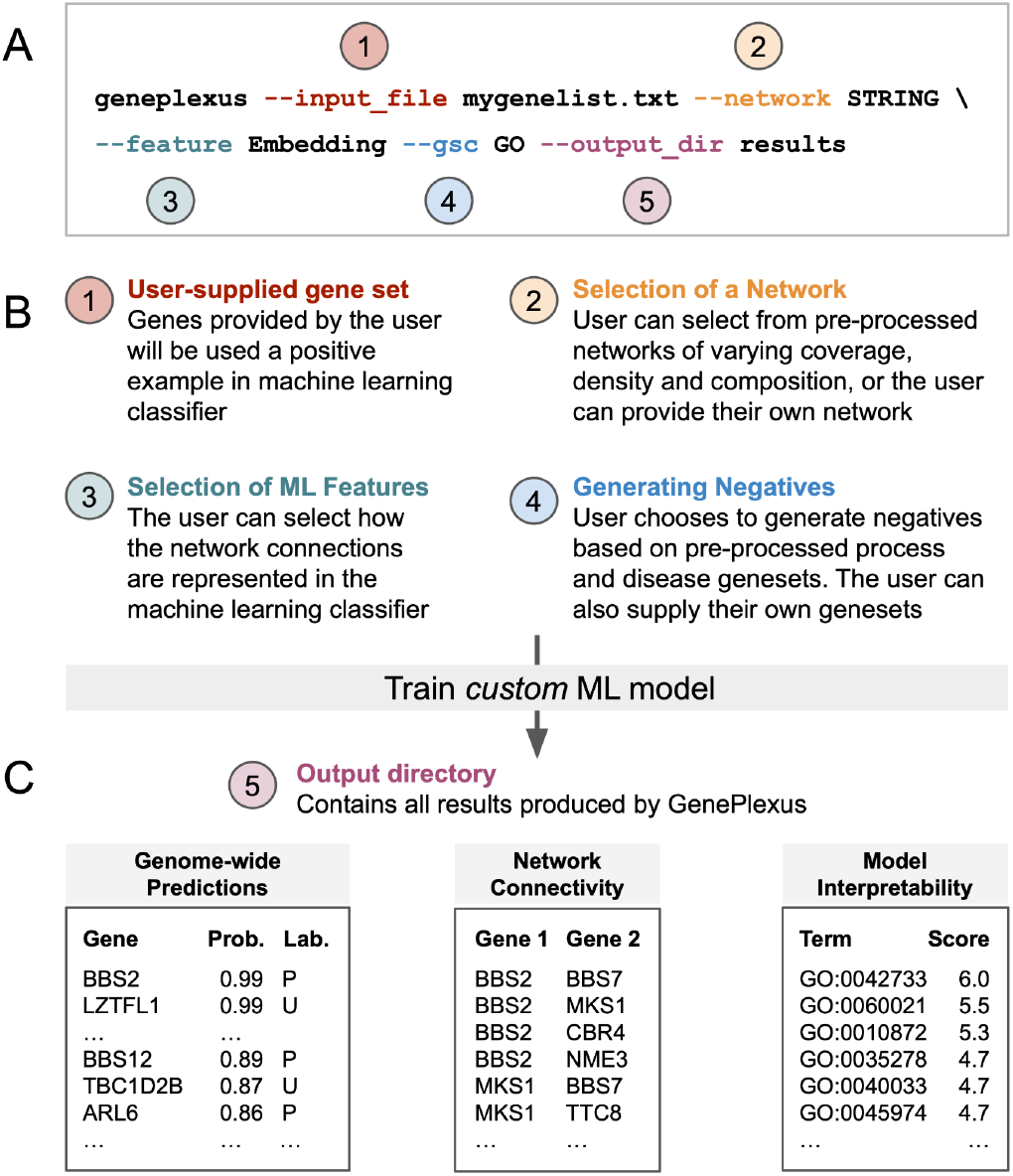
Running *PyGenePlexus* on the command line. (A) The *GenePlexus* model can be run with one simple command that (B) allows the user to select a number of different parameters and (C) obtain the results that are conveniently saved to the specified directory.

## Package overview

*PyGenePlexus* allows a user to input a set of genes and choose their desired network and its representation. *PyGenePlexus* then trains a custom ML model and returns the probability of how associated every gene in the network is to the user supplied gene set, along with the network connectivity of the top predicted genes. Additionally, the software provides an interpretation of the custom model by comparing it to thousands of models previously trained on gene sets from known biological processes and diseases. The following sections describe the different parts of the package.

### Downloading or supplying data

The *GenePlexus* method utilizes pre-processed information from genome-wide molecular networks and gene set collections from the Gene Ontology (GO) and DisGeNet. This data is archived on Zenodo (https://zenodo.org/record/6383205) and *PyGenePlexus* will automatically download the necessary data given the user input selections. Users can also supply their own networks and gene set collections to *PyGenePlexus*.

### Inputs

The user must first provide a set of genes, with valid ID types being Entrez, Symbols, Ensembl genes or Ensembl proteins [Fig 1B]. The user then chooses which molecular network to use and how that network should be represented in the ML model: as an adjacency matrix, an influence matrix, or a low-dimensional embedding of the network using *node2vec* (Grover and Leskovec, 2016; Liu and Krishnan, 2021). Finally, the user needs to choose which gene set collection (GO or DisGeNet) to use when determining the negative genes in the model.

## Results

Once the above parameters are chosen, a custom ML model is trained and the following results are returned [Fig. 1C]:

1. A prediction of how associated every gene in the network is to the input gene set.
2. The similarity of the model trained on the user-supplied gene set to thousands of models trained on gene sets from known pathways, processes and diseases.
3. The network connectivity of the top predicted genes.
4. The performance of the model based on *k*-fold cross-validation.

For more information on the pre-processed data, input choices, or results, see the package documentation.

### Example use case

The biological insights achievable using *PyGenePlexus* can be illustrated by considering the genes associated with *Bardet-Biedl syndrome 1* (BBS1) in the DisGeNet database (Supplemental File 1). The example below utilizes ‘BioGRID’ as the network, ‘embeddings’ as the feature representation, and ‘DisGeNet’ as the background for selecting negative genes. Examining the genome-wide predictions from *PyGenePlexus* (Supplemental Table 1) shows that the gene *LZTFL1* (*leucine zipper transcription factor like 1*) at rank 2 was originally not in the list of genes associated with the syndrome, and there is evidence that *LZTFL1* has a role in BBS1 (Marion *et al*., 2012). Comparison of the model trained on BBS1 genes to model trained on known disease gene set (Supplemental Table 2) shows that BBS1 model is highly similar to *Meckel syndrome* (both 8 and 1), which is a disease closely related to BBS1 (Karmous-Benailly *et al*., 2005; Forsythe and Beales, 2013). Comparison of the BBS1 model to models trained on gene sets from known biological process shows that the top ten results are terms relating to polydactylism, cholesterol and glycoside processes, and retina homeostasis, which relate to manifestations of BBS1 such as blindness, obesity, and having extra fingers or toes (Forsythe and Beales, 2013) (Supplemental Table 3).

## Discussion

*PyGenePlexus* is designed to be used by any researcher who wishes to gain insight about a gene set of interest using biological networks, even researchers with minimal programming knowledge. To help accomplish this, we provide extensive documentation of the package (https://pygeneplexus.readthedocs.io/en/main/). Additionally, *PyGenePlexus* can be run conveniently in two ways: pythonically through the class based method, or through a command line interface. Interacting directly with the Python code allows the user the ability to access all the functionalities of the package. The command line interface provides users who may not be familiar with Python an easier way to run the *PyGenePlexus* pipeline.

The *GenePlexus* method is also available through a well-documented, easy-to-use, interactive web-server (https://www.geneplexus.net/). *PyGenePlexus* offers some complementary functionalities not available on the web-server. First, *PyGenePlexus* allows a user to provide their own networks and gene set collections, which can be tailored to better fit the context in which their gene set was generated (e.g. through the use of tissue-specific gene interaction networks). Second, the local installation of *PyGenePlexus* allows a user to allocate any computational resources they have at hand to repeatedly run the pipeline, for example to predict on many gene sets or iterate through all the network-feature combinations on a given gene set. Thus, *PyGenePlexus* is a powerful, intuitive, well-documented tool that researchers with any level of programming ability can easily use to gain network-based biological insights into their gene sets of interest.

## Acknowledgment

We thank members of the Krishnan Lab for valuable discussions and feedback on the manuscript.

## Funding

This work was primarily supported by US National Institutes of Health (NIH) [grant R35 GM128765 to A.K.]; supported in part by National Institutes of Health [F32 F32GM134595 to C.A.M.].

## Conflict of interest

The authors have no conflict of interest to declare.

## Notes

https://github.com/krishnanlab/PyGenePlexus

https://pypi.org/project/geneplexus/

https://pygeneplexus.readthedocs.io/en/main/

https://zenodo.org/record/6383205

### Competing Interest Statement

The authors have declared no competing interest.

